# Where the patterns are: repetition-aware compression for colored de Bruijn graphs^⋆^

**DOI:** 10.1101/2024.07.09.602727

**Authors:** Alessio Campanelli, Giulio Ermanno Pibiri, Jason Fan, Rob Patro

**Affiliations:** DAIS, Ca’ Foscari University of Venice, Venice, Italy; ISTI-CNR, Pisa, Italy; Department of Computer Science, University of Maryland, College Park, MD 20440, USA

## Abstract

We describe lossless compressed data structures for the *colored* de Bruijn graph (or, c-dBG). Given a collection of reference sequences, a c-dBG can be essentially regarded as a map from *k*-mers to their *color sets*. The color set of a *k*-mer is the set of all identifiers, or *colors*, of the references that contain the *k*-mer. While these maps find countless applications in computational biology (e.g., basic query, reading mapping, abundance estimation, etc.), their memory usage represents a serious challenge for large-scale sequence indexing. Our solutions leverage on the intrinsic repetitiveness of the color sets when indexing large collections of related genomes. Hence, the described algorithms factorize the color sets into patterns that repeat across the entire collection and represent these patterns once, instead of redundantly replicating their representation as would happen if the sets were encoded as atomic lists of integers. Experimental results across a range of datasets and query workloads show that these representations substantially improve over the space effectiveness of the best previous solutions (sometimes, even dramatically, yielding indexes that are smaller by an order of magnitude). Despite the space reduction, these indexes only moderately impact the efficiency of the queries compared to the fastest indexes.

**Software:** The implementation of the indexes used for all experiments in this work is written in `C++17` and is available at https://github.com/jermp/fulgor.

## 1 Introduction

The colored de Bruijn graph (c-dBG) has become a fundamental tool used across several areas of genomics and pangenomics. For example, it has been widely adopted by methods that perform read mapping or alignment, specifically with respect to RNA-seq and metagenomic identification and abundance estimation [32,14,48,6,47,34,7,49]; among methods that perform homology assessment and mapping of genomes [38,39]; for a variety of different tasks in pangenome analysis [16,35,17,30,33], and for storage and compression of genomic data [46]. In most of these applications, a key requirement of the underlying representation of the c-dBG is to be able to determine — with efficiency being critical — the set of references (or “colors”) in which an individual *k*-mer appears. This set is named the *color set* of a *k*-mer^4^. These motivations bring us to the following problem formulation.

### Problem 1

(Colored *k*-mer indexing)

Let ℛ = {*R*_1_, …, *R*_*N*_ } be a collection of references. Each reference *R*_*i*_ is a string over the DNA alphabet *Σ* = {A, C, G, T}. We want to build a data structure (referred to as the *index* in the following) that allows us to retrieve the set ColorSet(*x*) = {*i*|*x* ∈ *R*_*i*_} as efficiently as possible for any *k*-mer *x* ∈ *Σ*^*k*^. If the *k*-mer *x* does not occur in any reference, we say that ColorSet(*x*) = ∅.

Of particular importance for biological analysis is the case where ℛ is a *pangenome*. Roughly speaking, a pangenome is a (large) set of genomes in a particular population, species or closely-related phylogenetic group. Pangenomes have revolutionized DNA analysis by providing a more comprehensive understanding of genetic diversity within a species [37,9]. Unlike traditional reference genomes, which represent a single individual or a small set of individuals, pangenomes incorporate genetic information from multiple individuals within a species or group. This approach is particularly valuable because it captures a wide range of genetic variations, including rare and unique sequences that may be absent from any particular reference genome.

### Contributions

The goal of this article is to propose compressed data structures to solve Problem 1 focusing on the specific, important, application scenario where ℛ is a pangenome. We note, however, that the approaches described herein are general, and we expect them to work well on any corpus of highly-related genomes, whether or not they formally constitute a pangenome. To best exploit the properties of Problem 1, we capitalize on recent indexing development for c-dBGs [21] that exploits an *order-preserving* dictionary of *k*-mers [42,43] to map *k*-mers to color sets in succinct space. This efficient mapping allows us to logically distinguish the problem of representing the dictionary of *k*-mers from that of representing the color sets. In this article, we therefore focus entirely on the latter problem.

We present two different solutions for the problem of representing the color sets in compressed space and explain how to combine them for even greater space reduction. These solutions are both based on the paradigm of *partitioning the color sets into patterns* that repeat across the whole collection of color sets. These patterns are encoded once in our data structures, thus avoiding unnecessary redundancy for their representation. This is in net contrast to previous methods in the literature, such as Fulgor [21] and Themisto [3], that represent distinct color sets only once, but which otherwise consider color sets as individual atomic integer lists and allow for repeated patterns within the color sets. Methods such as Mantis [5] and MetaGraph [28] do consider shared patterns and implement distinct but related forms of “differential” encoding, though the approaches are quite different from those introduced here and seem to require a larger sacrifice in query speed.

After covering preliminary concepts on efficient indexing of c-dBGs in Section 2, we review related work and the state of the art in Section 3. Section 4 describes our repetition-aware compression algorithms for color sets. In Section 5 we present a simple framework to build these compressed data structures. Section 6 presents experimental results to demonstrate that exploiting the repetitiveness of color patterns grants remarkably better compression effectiveness. As a result, the proposed solutions supersede all previous approaches in the literature, as they essentially combine the space effectiveness of the most compact methods with the query efficiency of the fastest solutions. This better efficiency comes at the expense of a slower construction algorithm. However, we do not regard this as a serious limiting aspect. We conclude in Section 7 where we also discuss some promising future work.

## 2 Preliminaries: modular indexing of colored de Bruijn graphs

In principle, Problem 1 could be solved using a classic data structure from information retrieval — the *inverted index* [45]. The inverted index is the backbone of virtually any retrieval system, mapping “terms” (e.g., words in natural language, or bag of words like *n*-grams, etc.) to the sorted lists of (the identifiers of) the documents that contain such terms. These sorted lists are called *inverted lists*. In the context of this problem, the indexed documents are the references {*R*_1_, …, *R*_*N*_ } in the collection ℛ and the terms of the inverted index are all the distinct *k*-mers of ℛ. Using the notation from Problem 1, it follows that ColorSet(*x*) is the inverted list of the term *x*. Let ℒ denote the inverted index for ℛ. The inverted index ℒ explicitly stores ColorSet(*x*) for each *k*-mer *x* ∈ ℛ. The goal is to implement the map *x* → ColorSet(*x*) as efficiently as possible in terms of both memory usage and query time. To this end, all the distinct *k*-mers of ℛ are stored in a dictionary 𝒟. Let *n* indicate the number of distinct *k*-mers in ℛ. These *k*-mers are stored losslessly in 𝒟. To be useful for this problem, the dictionary 𝒟 should be *associative*, that is: to implement the map *x* → ColorSet(*x*), 𝒟 is required to support the operation Lookup(*x*), which returns ⊥ if *k*-mer *x* ∈*/ 𝒟* or a *unique* integer identifier in [*n*] = {1, …, *n*} if *x* ∈ 𝒟.

Problem 1 can then be solved using these two data structures — 𝒟 and ℒ — thanks to the interplay between Lookup(*x*) and ColorSet(*x*): logically, the index stores the sets {ColorSet(*x*)}_*x*∈ℛ_ in some compressed form, *sorted by* the value returned by Lookup(*x*).

To exploit at best the potential of this modular decomposition into 𝒟 and ℒ, it is essential to rely on the specific properties of Problem 1. For example, we know that consecutive *k*-mers share (*k* − 1)-length overlaps; also, *k*-mers that co-occur in the same set of references have the same color set. A useful, standard, formalism that captures these properties is the so-called *colored de Bruijn graph* (c-dBG).

Let 𝒦 be the set of all the distinct *k*-mers of ℛ. The node-centric de Bruijn graph (dBG) of ℛ is a directed graph *G*(𝒦, *E*) whose nodes are the *k*-mers in 𝒦. There is an edge (*u, v*) ∈ *E* if the (*k* − 1)-length suffix of *u* equals the (*k* − 1)-length prefix of *v*. Note that the edge set *E* is implicitly defined by the set of nodes, and can therefore be omitted from subsequent definitions (there is a one-to-one correspondence between a dBG and a set of *k*-mers). We refer to *k*-mers and nodes in a dBG interchangeably. Likewise, a path (b) State-of-the-art index layout in a dBG spells the string obtained by concatenating together all the *k*-mers along the path, without repeating the shared (*k* − 1)-length overlaps. In particular, unary (i.e., non-branching) paths can be collapsed into single nodes spelling strings that are referred to as *unitigs*. Let 𝒰 = {*u*_1_, …, *u*_*m*_} be the set of unitigs of the graph. The dBG arising from this compaction step is called the *compacted* dBG, and indicated with *G*(𝒰).

The *colored* compacted dBG (c-dBG) is obtained by logically annotating each *k*-mer *x* with its color set, ColorSet(*x*). While different conventions have been adopted in the literature, here we assume that only non-branching paths with nodes having the *same* color set are collapsed into unitigs. The unitigs of the c-dBG we consider in this work have the following key properties.

### Property 1.

*Unitigs spell references in ℛ*. Each distinct *k*-mer of ℛ appears once, as sub-string of some unitig of the c-dBG. By construction, each reference *R*_*i*_ ∈ ℛ can be spelled out by some *tiling* of the unitigs — an ordered sequence of unitig occurrences that, when glued together (accounting for (*k*-1)-symbol overlap and orientation), spell exactly *R*_*i*_ [20]. Joining together *k*-mers into unitigs reduces their storage requirements and accelerates looking up *k*-mers in consecutive order [42].

### Property 2.

*Unitigs are monochromatic*. The *k*-mers belonging to the same unitig *u*_*i*_ all have the same color set. We write *x* ∈ *u*_*i*_ to indicate that *k*-mer *x* is a sub-string of the unitig *u*_*i*_. Thus, we shall use ColorSet(*u*_*i*_) to denote the color set of each *k*-mer *x* ∈ *u*_*i*_. This suggests that a single color set should be represented *per unitig*, rather than for each *k*-mer.

### Property 3.

*Unitigs co-occur*. Distinct unitigs often have the *same* color set, i.e., they co-occur in the same set of references, because they derive from conserved sequences in indexed references that are longer than the unitigs themselves. Hereafter, we indicate with *z* the number of distinct colors set 𝒞 = {*C*_1_, …, *C*_*z*_}. Note that *z* ≤ *m* and that, in practice, there are almost always *many more* unitigs than there are distinct color sets.

Figure 1a illustrates an example c-dBG with these properties. In the following sections, we refer to a compacted c-dBG as *G*(𝒰, *𝒞*).

**Fig. 1:**
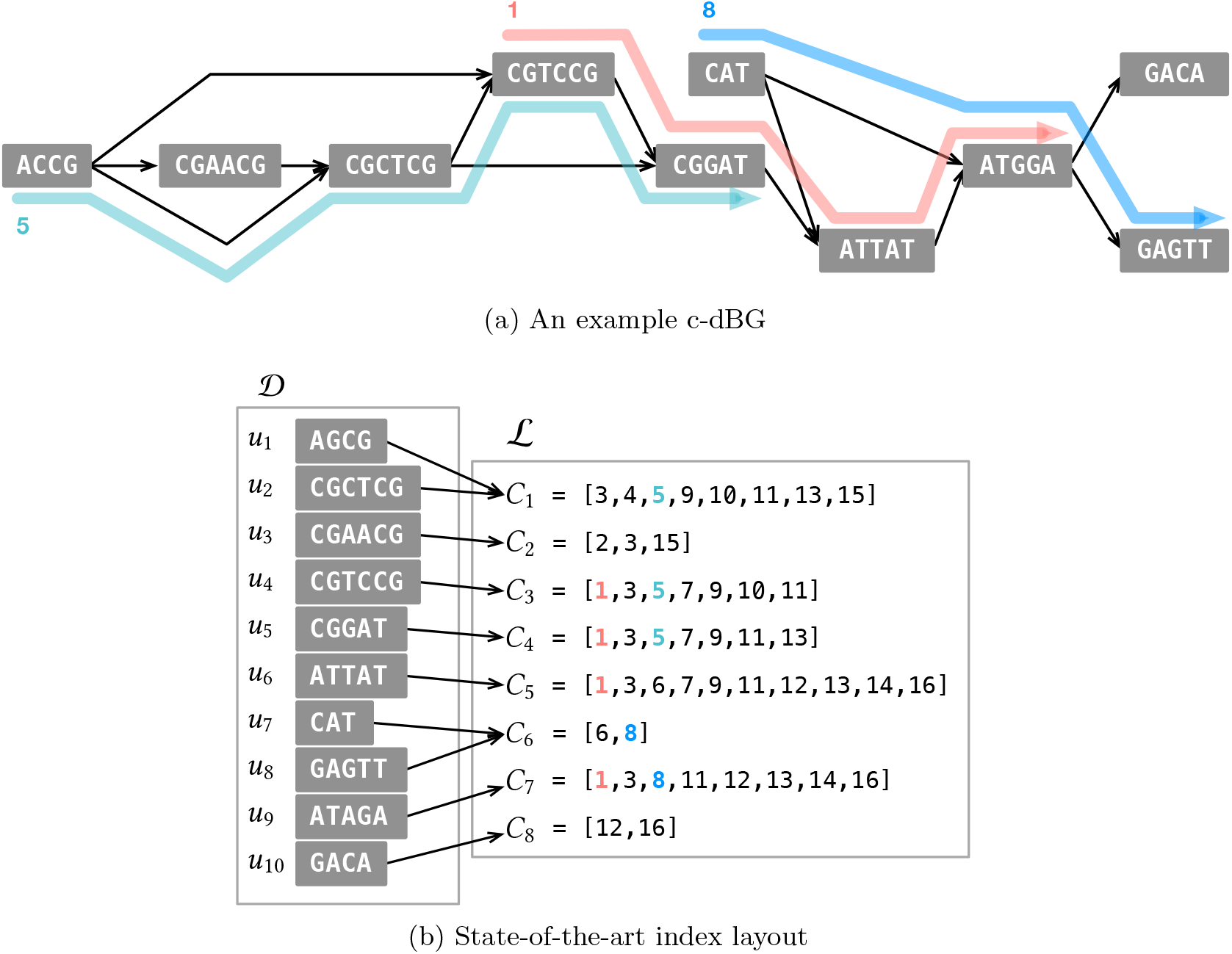
In panel (a), an example c-dBG for *k* = 3 with three colors (1, 3, and 8) highlighted. (In the figure, a *k*-mer and its reverse complement are considered as different *k*-mer for ease of illustration. In practice, these are considered identical.) The unitigs of the graph are colored according to the set of references they appear in. In panel (b), we schematically illustrate the state-of-the-art index layout (as used in the Fulgor index [21]) assuming the c-dBG was built for *N* = 16 references, showing the modular composition of a *k*-mer dictionary, 𝒟, and an inverted index, ℒ. Note that unitigs are stored in 𝒟 in *color set order* (i.e., unitigs mapping to the same color set are placed consecutively in the dictionary), thereby allowing a very efficient mapping of *k*-mers to their distinct colors.

The modular indexing layout described in this section allows us to logically divide Problem 1 into two sub-problems: (1) that of representing *k*-mer sets under Lookup queries, and (2) that of compressing the collection of color sets 𝒞 = {*C*_1_, …, *C*_*z*_}. This work specifically targets the latter problem. Our solutions are presented in Section 4.

## 3 Related work

The data structures proposed in the literature to represent c-dBGs, and that fall under the “color-aggregative” classification [36], all provide different implementations of the modular indexing framework as described in Section 2. As such, they require two data structures: (1) a *k*-mer dictionary and (2) an inverted index. The methods reviewed below differ in the choice of the dictionary and the compression scheme for the inverted index.

For example, Themisto [3] makes use of the *spectral* BWT (SBWT) data structure [2] for its *k*-mer dictionary and an inverted index compressed with different strategies based on the sparseness of the color sets (i.e., the ratio |*C*_*i*_|*/N*). Mantis [40,5,4] makes use of the counting quotient filter [41] for the dictionary data structure, and in its most space-efficient variant represents the color sets by deduplicating them and expressing them differentially as edits performed along the branches of an approximate minimum spanning tree over the color sets. MetaGraph [28] uses the BOSS [13] data structure for the dictionary and exposes several general schemes to compress metadata associated with *k*-mers [28,29], which essentially constitute an inverted index. Bifrost [24], instead, uses a (dynamic) hash table to store the set of unitigs and an inverted index compressed with Roaring bitmaps [31]. The compact bit-sliced (COBS) index [10] can be considered as an *approximate* c-dBG in that the ColorSet(*x*) might contain false positives, i.e., spurious reference identifiers (but never false negatives). This is a consequence of building a Bloom filter [12] for each reference, filled with all the *k*-mers in the reference. The Bloom filter matrix is stored in an inverted manner, and represents a collection of approximate color sets. Being approximate, this method completely avoids the space consumption of the exact *k*-mer dictionary and the space is all due to the approximate color sets.

However, none of these solutions simultaneously exploit all three unitig properties listed in Section 2 to achieve faster query time and better space effectiveness. More specifically, Themisto disregards Property 1 as a direct consequence of using the SBWT data structure that internally arranges the *k*-mers in *colexicographic* order, and not in their order of appearance in the unitigs. This translates to an overhead of log_2_(*z*) bits^5^ per *k*-mer to associate a *k*-mer to its color set (we recall that *z* indicates the number of distinct color sets in our discussion). This consideration is also valid of the BOSS data structure, hence for MetaGraph and the counting quotient filter data structure and hence for Mantis. Themisto exploits Property 3 instead, by compressing only the set of the *distinct* color sets. Alanko et al. also describe how it is possible in Themisto to reduce the space for the mapping from *k*-mers to colors by spending log_2_(*z*) bits for only some *k*-mers (called *core k*-mers in their work), while instead using 1 + *o*(1) bits per *k*-mer for all the others. However, this still requires dedicated storage *per*-*k*-mer, thus failing to exploit Property 2. COBS does not exploit any specific property, instead: unitigs are broken into their constituent *k*-mers and indexed separately; looking up consecutive *k*-mers (most likely part of the same unitig) has no locality of reference due to Bloom filter lookups; color sets are stored approximately and partitioned into shards, so that a ColorSet(*x*) query has to combine several partial results together.

### State of the art

To the best of our knowledge, the only solution that exploits *all* the three properties is the recently-introduced Fulgor index [21]. The solution adopted by Fulgor is to first map *k*-mers to unitigs using 𝒟, and then succinctly map unitigs to their color sets. By *composing* these two mappings, Fulgor obtains an efficient map directly from *k*-mers to their associated color sets. Refer to Figure 1b for an illustration of this index. The composition is made possible by leveraging the *order-preserving* property of its dictionary — SSHash [42,43] — which explicitly stores the set of unitigs in *any* desired order. This property has some important implications. First, looking up consecutive *k*-mers is cache-efficient since unitigs are stored contiguously in memory as sequences of 2-bit characters. Second, if *k*-mer *x* occurs in unitig *u*_*i*_, the Lookup(*x*) query of SSHash can efficiently determine the unitig identifier *i*, allowing to map *k*-mers to unitigs. Third, if unitigs are sorted in *color set order*, so that unitigs having the same color set are consecutive, then mapping a unitig identifier *i* to its color set identifier, ColorSet-ID(*i*), can be implemented in as little as 1 + *o*(1) bits *per unitig* and in constant time. This is achieved using a succinct Rank query, as explained in Fact 1 below.

Given a bitvector *b*[1..*u*], we let Rank_1_(*b, i*) be the number of bits set in *b*[1..*i*), for any 1 ≤ *i* ≤ *u*. It is possible to answer Rank queries in *O*(1) using just *o*(*u*) bits on top of the plain bitvector *b* [26].

### Fact 1

Consider a sorted collection of *u* items where *f* ≤ *u* are distinct. Call *v*_*i*_ the *i*-th item in the sorted order. Given an index 1 ≤ *i* ≤ *u*, we can determine *v*_*i*_ in 𝒪 (1) without storing all the *u* items explicitly, as follows. We keep the *f* distinct items in an array *A*[1..*f*] and build a bitvector *b*[1..*u*] where: *b*[*i*] = 1 if and only if *v*_*i*_≠ *v*_*i*+1_ for any 1 ≤ *i < u* and *b*[*u*] = 1. (Note that *b* has exactly *f* bits set.) With *b* and *A*, we recover *v*_*i*_ as *A*[*p*] where *p* = Rank_1_(*b, i*) + 1. Apart from the space of *A* which is at most that of the original collection because *f* ≤ *u*, this solution spends 1 + *o*(1) bits per item.

We used Fact 1 to succinctly map unitigs to their color sets in Fulgor by letting *v*_*i*_ = ColorSet-ID(*i*) and the array *A* be C. We will use Fact 1 in this article as well.

Lastly, the color sets 𝒞 = {*C*_1_, …, *C*_*z*_} themselves are compressed in Fulgor using three different encodings based on the ratio |*C*_*i*_|*/N*. If |*C*_*i*_|*/N <* 1*/*4, then the set *C*_*i*_ is considered *sparse* and the differences between consecutive integers are computed and encoded with Elias’s δ code [19]. If, instead, |*C*_*i*_|*/N >* 3*/*4, the set is considered *very dense* and the complementary set [*N*] ∩ *C*_*i*_ is compressed using the method explained before. Lastly, if 1*/*4 ≤ |*C*_*i*_|*/N* ≤ 3*/*4, the set is *dense* and represented as a bitvector of *N* bits where the *j*-th bit is set if color *j* ∈ *C*_*i*_. Note that each set *C*_*i*_ is thus encoded *individually* from the other sets in C.

## 4 Repetition-aware compression for colored de Bruijn graphs

When indexing large pangenomes, the space taken by the color sets 𝒞 = {*C*_1_, …, *C*_*z*_}, even in compressed format, dominates the whole index space [21,24,3]. Efforts toward improving the memory usage of c-dBGs should therefore be spent in devising better compression algorithms for the color sets. In this work, we focus on exploiting the following crucial property that can enable substantially better compression effectiveness: *The genomes in a pangenome are very similar*. This, in turn, implies that the *color sets are also very similar* (albeit distinct).

By “similar” sets we mean that they share many (potentially, very long) identical integer sub-sequences. This property is not exploited if each set *C*_*i*_ is compressed *individually* from the other sets. For example, if set *C*_*i*_ shares a long sub-sequence with *C*_*j*_, this sub-sequence is actually represented *twice* in the index, which wastes space. We refer to such shared sub-sequences as “patterns” in the following. Consider Figure 2 for some examples. The pattern [3, 5, 9, 11] repeats in *C*_1_, *C*_3_, and *C*_4_, hence it is represented redundantly three times. The longer pattern [1, 3, 11, 12, 13, 14, 16] repeats twice instead. These examples are instrumentally simple; yet, they suggest that the identification of such patterns across a large collection, as well as the design of an effective compression mechanism for these patterns, is not easy. A further complicating matter is that these patterns can repeat in *many* sets (not just two or three), hence increasing the pangenome redundancy and aggravating the memory usage of an index that encodes them redundantly in the many sets where they appear.

**Fig. 2:**
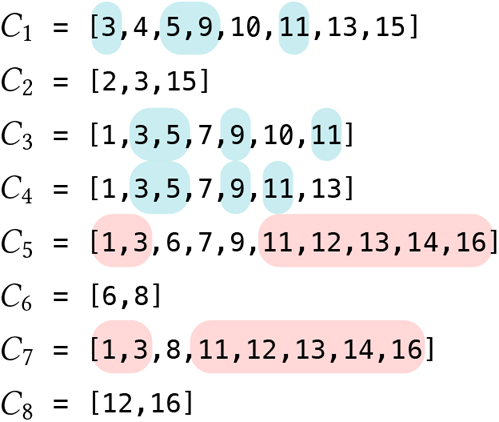
The color sets from Figure 1b where we highlight some “patterns” (i.e., repetitive sub-sequences) that repeat across different sets, like [3, 5, 9, 11] and [1, 3, 11, 12, 13, 14, 16].

To address this issue, we propose here two solutions based on *partitioning* the color sets with the intent of factoring out repetitive patterns. Encoding such repetitive patterns *once* clearly reduces the amount of redundancy in the representation, which improves the space of the data structures. We explore the effectiveness of two different partitioning paradigms which, for ease of visualization, we refer to as *horizontal* partitioning (Section 4.1) and *vertical* partitioning (Section 4.2). Figure 3 shows an example of partitioning that we will refer to in the following sections. In Section 4.3 we argue that these two partitioning paradigms can be combined to improve compression even further.

**Fig. 3:**
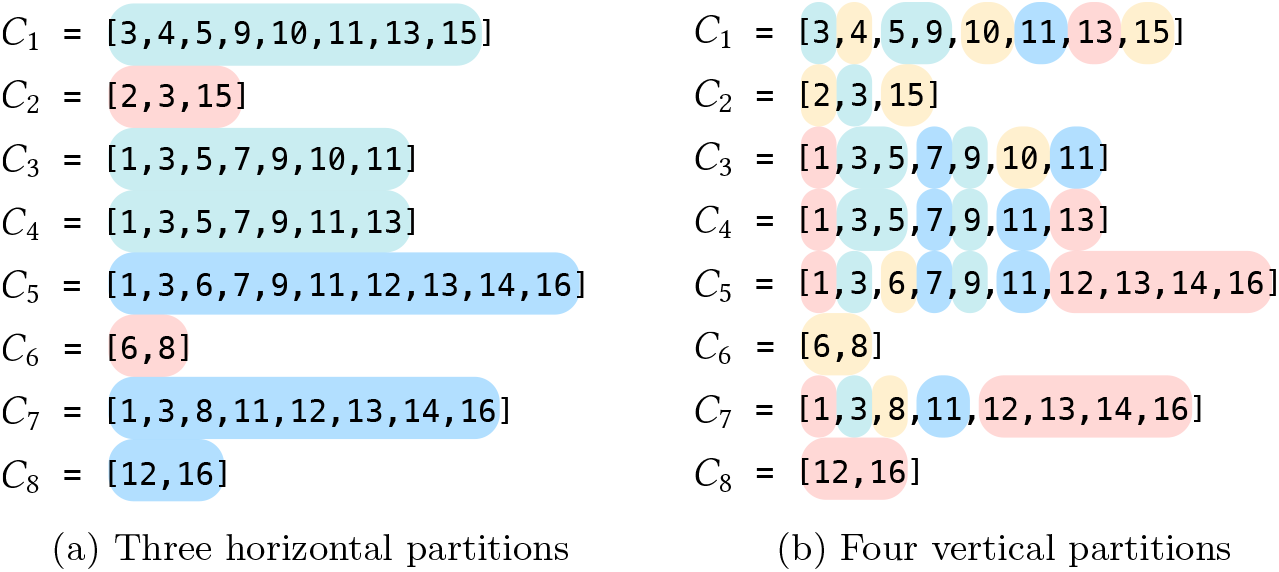
The color sets from Figure 1b, respectively partitioned horizontally (a) and vertically (b) Intuitively, partitions of “similar” *rows* are created by horizontal partitioning; vice versa, partitions of similar *columns* are created by vertical partitioning.

Before presenting the details of our solutions, we first establish the following fact. Given an integer *q* ≥ 1, let 𝒩 = {𝒩_1_, …, *𝒩*_*r*_} be a partition of [*q*] = {1, …, *q*} of size *r* ≥ 1. Let an order between the elements {*e*_*ij*_} of each N_*i*_ be fixed (for example, by sorting the elements {*e*_*ij*_} in increasing order).

**Fact 2**. Any 𝒩 induces a permutation *π* : [*q*] → [*q*], defined as *π*(*e*_*ij*_) := *j* + *B*_*i*−1_ where 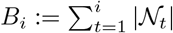 for *i >* 0 and *B*_0_ := 0, for *i* = 1, …, *r* and *j* = 1, …, |𝒩_*i*_|.

Consider the following example for *q* = 10 and *r* = 3. Suppose 𝒩_1_ = {3, 7, 9}, 𝒩_2_ = {1, 4}, and 𝒩_3_ = {2, 5, 6, 8, 10}. The boundaries *B*_*i*_ are therefore *B*_0_ = 0, *B*_1_ = 3, *B*_2_ = 5, and *B*_3_ = 10. The induced permutation *π* can be visualized by concatenating the sets 𝒩_*i*_ from *i* = 1 to 3 and assigning “new” identifiers, from 1 to *q*, in this concatenated order:

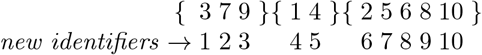

which results in *π*(3) = 1, *π*(7) = 2, *π*(9) = 3, etc., that is *π* = [4, 6, 1, 5, 7, 8, 2, 9, 3, 10].

### 4.1 Horizontal partitioning: representative and differential color sets

The first solution we present explores horizontal partitioning. In short, the general idea is that of partitioning the sets in C into groups of similar sets (see Figure 3a for an example); then, for each group, a *representative* color set is built and all sets in the group are encoded via a *differential* set with respect to the representative one.

**Definition**. Let 𝒩 be a partition of [*z*], of size *r* ≥ 1, and let *π* be its corresponding permutation as established by Fact 2. For each 𝒩_*i*_, we build a set *A*_*i*_ in some way that we will explain shortly and represent the set *C*_*j*_ as (*A*_*i*_ Δ*C*_*j*_) for all *j* ∈ 𝒩_*i*_. Notation (*X* Δ*Y*) stands for the symmetric difference between the sets *X* and *Y*, and it is (*X* ∪ *Y*) \ (*X* ∩ *Y*). The idea is that the set *A*_*i*_ should include the most repetitive patterns that occur in the sets of partition 𝒩_*i*_, so that each difference (*A*_*i*_ Δ*C*_*j*_) is *small*. Since *A*_*i*_ should capture the repetitiveness of 𝒩_*i*_, it is named the *representative* color set of 𝒩_*i*_. The symmetric difference (*A*_*i*_ Δ*C*_*j*_) is instead called the *differential* color set of *C*_*j*_ with respect to *A*_*i*_. We indicate with 𝒜 the set of all representative color sets. Note that |𝒜| = *r* as we have one representative per partition. Similarly, we indicate with Δ the set of all differential color sets, where |Δ| = *z* as we have one differential color set for each original set in 𝒞.

Both sets, 𝒜 and Δ, are therefore made up of sequences of increasing integers that can be compressed effectively using a plethora of different methods [45]. We discuss implementation details in Section 6.

Compared to a c-dBG *G*(𝒰, 𝒞), its differential-colored c-dBG (or, Dfc-dBG) variant is the graph *G*(𝒰, *𝒩, π, 𝒜*, Δ) where the set of nodes, 𝒰, is the same as that of *G* but the sets in C are factorized into 𝒜 and Δ. We now illustrate an example of how the sets in C can be modeled via 𝒜 and Δ.

**Example**. Let us consider the *r* = 3 partitions from Figure 3a for the *z* = 8 color sets as highlighted by different shades, i.e., 𝒩_1_ = {1, 3, 4}, 𝒩_2_ = {2, 6}, and 𝒩_3_ = {5, 7, 8}. Assume the following representative color sets, *A*_1_, *A*_2_, and *A*_3_, are built.

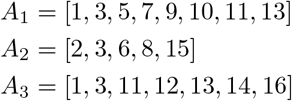

we then obtain the following differential color sets.

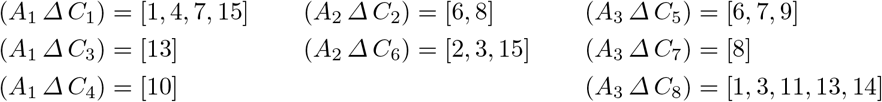

Note how the pattern [3, 5, 9, 11] shared by all the three sets *C*_1_, *C*_3_, and *C*_4_ is now encoded once in the representative set *A*_1_ and *implicitly* encoded in each differential set (*A*_1_ Δ*C*_1_), (*A*_1_ Δ*C*_3_), and (*A*_1_ Δ*C*_4_). Consider the pattern [1, 3, 11, 12, 13, 14, 16] shared by *C*_5_ and *C*_7_ for another example (in this case, the pattern is exactly equal to the representative set *A*_3_). If, instead, the representative color set does not include patterns that appear in sufficiently many sets in its partition, then the representation is wasteful. This is the case, in this example, for the set *A*_2_. In the worst case, when the intersection between *A*_*i*_ and *C*_*j*_ is empty, then (*A*_*i*_ Δ*C*_*j*_) = *A*_*i*_ ∪ *C*_*j*_, which is more than storing the set *C*_*j*_ itself.

To conclude, Figure 4 shows all the components of the Dfc-dBG from this example. Importantly, note that the order of the sets {*C*_1_, …, *C*_8_} has been permuted according to *π* to efficiently determine the partition of any given color set identifier as to perform decoding of the set (see the following discussion for more details).

**Fig. 4:**
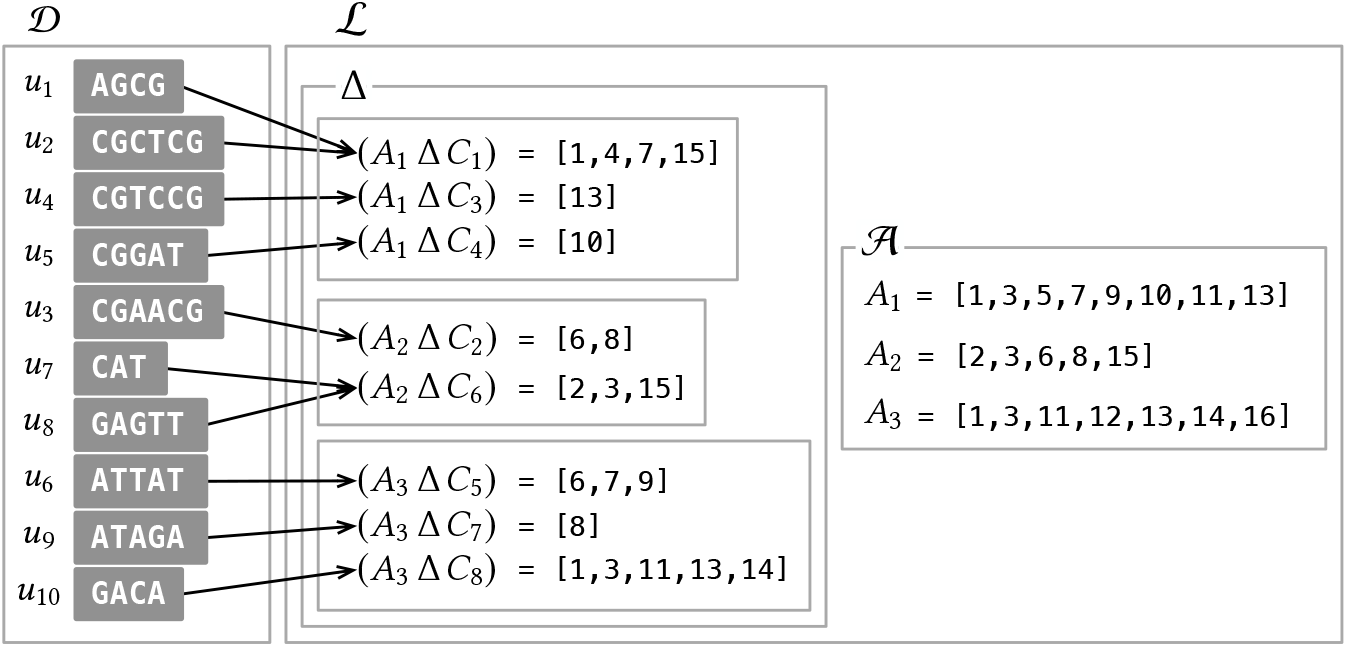
The Dfc-dBG layout derived from the color sets in Figure 1b. Importantly, note that color sets are stored in the order given by the permutation *π* induced by the horizontal partitioning. This allows mapping a color set identifier to its partition in very succinct space. Note that also the unitigs in 𝒟 are re-sorted following the permuted order of the color sets.

### Discussion

We now discuss the salient features of the introduced Dfc-dBG. All in all, the Dfc-dBG saves space compared to the original c-dBG with only a minor slowdown when performing a ColorSet(*x*) query.

1. As already explained, if the representative color sets include the most repetitive patterns in their respective partitions, then the size of differential color sets is expected to be small, thereby reducing the amount of integers that are represented in the index. This clearly saves space compared to *G*(𝒰, *𝒞*) because the memory usage of the index is proportional to the number of integers being encoded.
2. The order in which the differential color sets are stored in the index is not necessarily 1, …, *z*, as illustrated in the layout from Figure 1b. The order is instead obtained by applying *π* to [*z*]. In our example from Figure 4, the permutation *π* is [1, 4, 2, 3, 6, 5, 7, 8], hence the color sets *C*_1_, …, *C*_8_ are stored in the order *C*_1_, *C*_3_, *C*_4_, *C*_2_, *C*_6_, *C*_5_, *C*_7_, *C*_8_, given that *π*(1) = 1, *π*(3) = 2, *π*(4) = 3, etc. So, color sets belonging to the same partition are placed consecutively by *π*. This clearly permits to determine the partition a color belongs to in an efficient way, e.g., using Fact 1. Specifically, we use a binary vector *b*[1..*z*] and let *b*[*i*] = 1 if and only if *i* is the position of the last color set of a partition. Continuing our example, we have

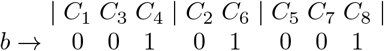

where *b*[3] = *b*[5] = *b*[8] = 1 because the third, the fifth, and the eighth color set in the permuted order are the last in their respective partitions. To conclude, using 1 + *o*(1) extra bits per color, we can compute the partition 1 ≤ *p* ≤ *r* of color *C*_*j*_ as *p* = Rank_1_(*b, π*(*j*)) + 1 in constant time as per Fact 1.
3. Lastly, it is efficient to decode *C*_*j*_ from (*A*_*i*_ Δ*C*_*j*_). By definition of symmetric set difference, it follows that *C*_*j*_ = (*A*_*i*_ Δ (*A*_*i*_ Δ*C*_*j*_)). Hence decoding *C*_*j*_ corresponds to computing the symmetric difference between the representative color set and the differential color set. This can be simply implemented in time linear in the size of the two sets. Note that |*C*_*j*_| ≤ |*A*_*i*_|+|(*A*_*i*_ Δ*C*_*j*_)| (with equality holding only when *A*_*i*_ = ∅), hence decoding takes more time than just scanning the original color *C*_*j*_. This imposes some overhead compared to any representation encoding each set individually. However, as we are going to see in Section 6, this overhead is not much because decoding is cache-friendly.

#### The optimization problem

The effectiveness of the Dfc-dBG evidently depends on the choice of the partition 𝒩, i.e., what color sets to cluster together, and the choice of the representative for each partition 𝒩_*i*_. Intuitively, one would like to group similar color sets together and let their shared patterns be included in their representative color set. On one hand, the number of partitions, *r*, should not be too large to amortize the cost of the representative color sets; on the other hand, smaller partitions better highlight the repetitiveness of the patterns in the collection. In the following, let Cost(*L*) be the encoding cost of the sorted list *L* according to some encoding method. We can formally state the optimization problem faced by the Dfc-dBG representation as follows. We call it the *minimum-cost partition arrangement* (MPA) problem for Dfc-dBG.

##### Problem 2

(MPA for Dfc-dBG).

Let *G*(𝒰, *𝒞*) be the compacted c-dBG built from the reference collection ℛ = {*R*_1_, …, *R*_*N*_ }, where |*𝒞*| = *z*. Determine the partition 𝒩 = {𝒩_1_, …, *𝒩*_*r*_} of [*z*] for some *r* ≥ 1 and the sets *A*_1_, …, *A*_*r*_ such that 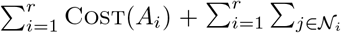 Cost(*A* Δ*C*) is minimum.

We suspect this problem is hard depending on the chosen encoding method. Instead, we prove the following.

##### Theorem 1.

Given a partition 𝒩 = {𝒩_1_, …, *𝒩*_*r*_} of [*z*] of size *r* ≥ 1, let Cost(*L*) = |*L*| and

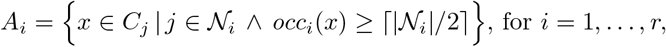

where 1 ≤ *occ*_*i*_(*x*) ≤ |𝒩_*i*_| denotes the number of occurrences of the integer *x* in the color sets of 𝒩_*i*_. Then the cost 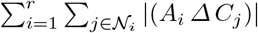 is minimum.

The rationale behind the choice Cost(*L*) = |*L*| is to minimize the number of integers being encoded in the differential color sets, which is the most expensive component in the Dfc-dBG.

*Proof*. By contradiction. Assume *A*_*i*_ is optimal and there exists an integer *x* ∈ *A*_*i*_ such that *occ*_*i*_(*x*) *<* ⌈|𝒩_*i*_|*/*2⌉. This means that there are |𝒩_*i*_|− *occ*_*i*_(*x*) differential color sets containing *x*. Let 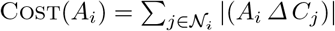. We can therefore remove *x* from *A*_*i*_ to obtain a new solution 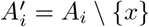 such that

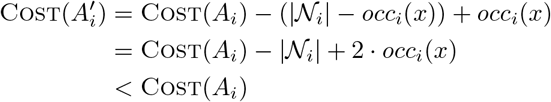

where the last inequality holds because *occ*_*i*_(*x*) *<* ⌈|𝒩_*i*_|*/*2⌉. Solution 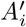 has therefore a lower cost than *A*_*i*_. This contradicts the initial assumption that *A*_*i*_ is optimal. ■

The representative color sets in Figure 4 and in all previous examples are built with the strategy described in Theorem 1. Consider, for example, the color sets in the partition 𝒩_1_ = {*C*_1_, *C*_3_, *C*_4_} from Figure 3a. In this case, |𝒩_1_| = 3 and ⌈|𝒩_1_|*/*2⌉ = 2, so any integer appearing at least twice among those in *C*_1_, *C*_2_, and *C*_3_, is included in *A*_1_. Specifically, we have *occ*_1_(1) = 2, *occ*_1_(3) = 3, *occ*_1_(4) = 1, *occ*_1_(5) = 3, etc. It is easy to see that the only two integers that cannot make it into *A*_1_ are 4 and 15 since *occ*_1_(4) = *occ*(15) = 1 *<* 2. In conclusion, we have *A*_1_ = [1, 3, 5, 7, 9, 10, 11, 13].

The quantities *occ*_*i*_(*x*) can be computed by iterating through all color sets in C once and, given that there are at most *N* distinct integers in each partition 𝒩_*i*_, the sets *A*_1_, …, *A*_*r*_ are computed in a total of *O*(Σ _*i*_ |*C*_*i*_| + *r* · *N* log *N*) time.

### 4.2 Vertical partitioning: partial and meta color sets

In this section we present a second solution, based on vertical partitioning (see Figure 3b for an example). Now, each color set in C is spelled by a list of references, or *meta colors*, to smaller repetitive patterns that we call *partial* color sets.

**Definition**. Let 𝒩 = {𝒩_1_, …, *𝒩*_*r*_} be a partition of [*N*], of size *r* ≥ 1, and refer to *π* as its induced permutation as per Fact 2. We assume from now on that the *N* reference identifiers and the integers in the sets of 𝒞 have been permuted according to *π*. After the permutation, 𝒩 determines a partition of ℛ into *r* disjoint sets:

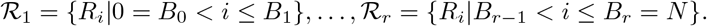

Let 𝒫_*i*_ be the set

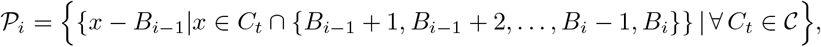

for *i* = 1, …, *r*. The elements {*P*_*ij*_} of the set 𝒫_*i*_ are the *partial color sets* induced by the partition 𝒩_*i*_. We indicate with 𝒫 = {𝒫_1_, …, *𝒫*_*r*_} the set of all partial color sets. In simple words, 𝒫_*i*_ is the set obtained by considering the distinct color sets *only* for the references in the *i*-th partition ℛ_*i*_, noting that — by construction — they comprise integers *x* such that *B*_*i*−1_ *< x* ≤ *B*_*i*_. The idea is that the set 𝒫 = {𝒫_1_, …, *𝒫*_*r*_} forms a dictionary of subsequences (the partial color sets) that spell the original color sets 𝒞 = {*C*_1_, …, *C*_*z*_}. Let us now formally define this spelling.

Let *C*_*t*_ ∈ 𝒞 be a color set. A *meta color* is an integer pair (*i, j*) indicating the sub-list *L* := *C*_*t*_[*b*..*b* + |*P*_*ij*_|] if there exists 0 *< b* ≤ |*C*_*t*_| − |*P*_*ij*_| such that *L*[*l*] = *P*_*ij*_[*l*] + *B*_*i*−1_, for *l* = 1, …, |*P*_*ij*_|. It follows that *C*_*t*_ can be modeled as a list *M*_*t*_ of at most *r* meta colors. We indicate with ℳ = {*M*_1_, …, *M*_*z*_} the set of all meta color sets.

We have therefore obtained a representation of 𝒞 that consists of the sets ℳ and 𝒫. These sets are, again, made up of integer sequences that can be further compressed. For the experiments in Section 6, we will choose a suitable compression methods for these sets.

Given *G*(𝒰, 𝒞), the meta-colored c-dBG (or, Mac-dBG) is the graph *G*(𝒰, *𝒩, π, 𝒫, ℳ*) where the set of nodes, 𝒰, is the same as that of *G* but the color sets in C are represented with the partial color sets 𝒫 and the meta color sets ℳ. We now illustrate an example.

**Example**. Let us consider the *z* = 8 color sets from Figure 1b, for *N* = 16. Let *r* = 4 and 𝒩_1_ = {1, 12, 13, 14, 16}, 𝒩_2_ = {3, 5, 9}, 𝒩_3_ = {7, 11}, 𝒩_4_ = {2, 4, 6, 8, 10, 15}, assuming we use the natural order between the integers to determine an order between the elements of each 𝒩_*i*_. Thus, we have *B*_1_ = 5, *B*_2_ = 8, *B*_3_ = 10, and *B*_4_ = 16. The corresponding permutation *π* is therefore [1, 11, 6, 12, 7, 13, 9, 14, 8, 15, 10, 2, 3, 4, 16, 5]. Now we apply the permutation *π* to each color set, obtaining the following permuted color sets (vertical bars represent the partial color set boundaries *B*_1_, *B*_2_, *B*_3_).

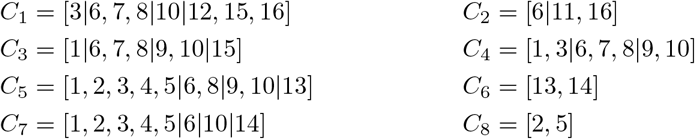

For example, set *C*_1_, that before was [3, 4, 5, 9, 10, 11, 13, 15] (see Figure 1b), now is

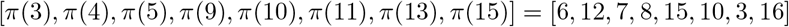

or [3, 6, 7, 8, 10, 12, 15, 16] once sorted. The partial color sets are the distinct sub-sequences in each partition of the permuted color sets. For example, 𝒫_1_ is the set of the distinct subsequences in partition 1, i.e., those comprising the integers *x* such that 0 *< x* ≤ *B*_1_ = 5. Hence, we have five distinct partial color sets in partition 1, and these are [3], [1], [1, 3], [1, 2, 3, 4, 5], and [2, 5]. Importantly, note that from the integers in the partial color sets of partition *i >* 1 we can subtract the lower bound *B*_*i*−1_. For example, we can subtract *B*_1_ = 5 from the integers of [6, 7, 8], that is the partial color set used in the partition 2 of *C*_1_, to obtain [1, 2, 3]. Overall, we thus obtain that 𝒫 comprises four partial color sets, as shown in Figure 5. The figure also shows the rendering of the color sets 𝒞 = {*C*_1_, …, *C* _8_} via meta color lists, i.e., how each color can be spelled by a proper concatenation of partial color sets.

**Fig. 5:**
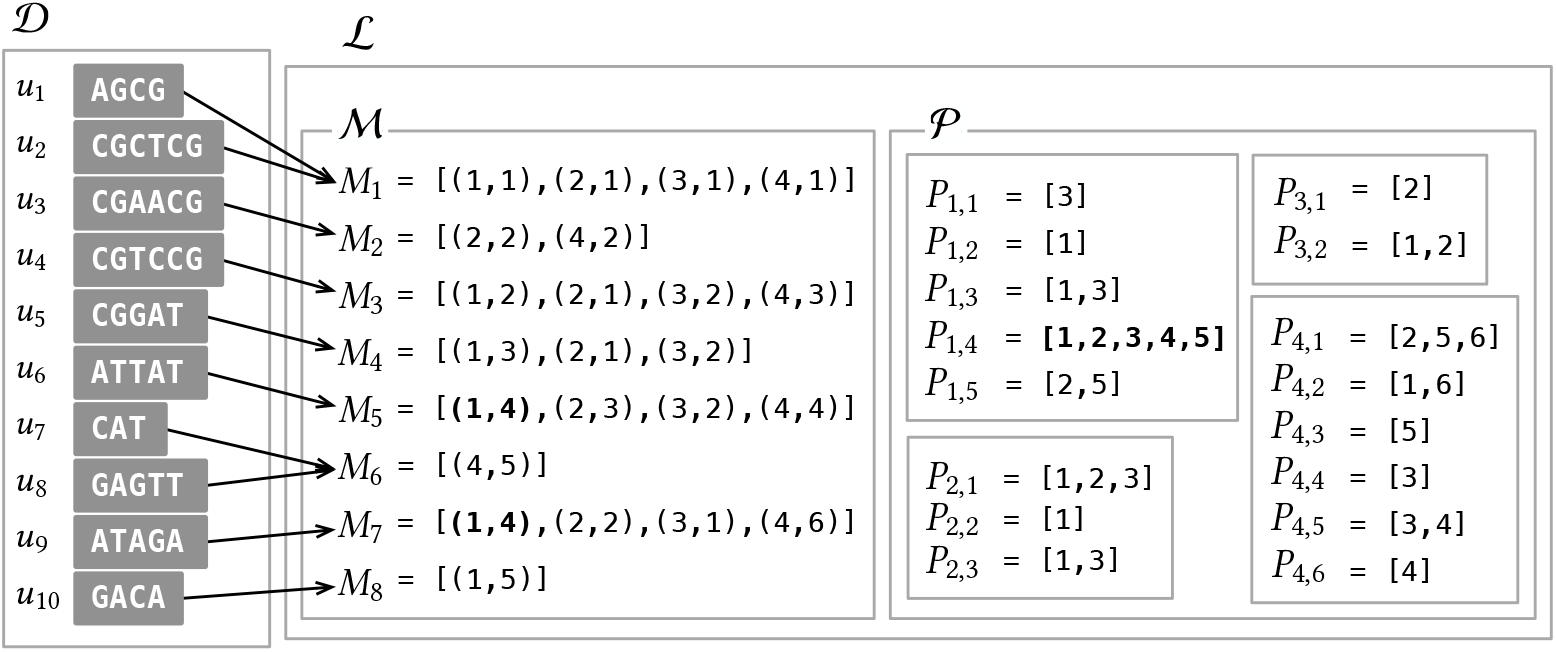
The Mac-dBG layout derived from the color sets in Figure 1b. Note that the partial color set *P*_1,4_ = [1, 2, 3, 4, 5] shared between *C*_5_ and *C*_7_ is now represented *once* as a direct consequence of partitioning, and indicated with the meta color (1, 4) instead of replicating the five integers it contains in both *C*_5_ and *C*_7_. The same consideration applies to other shared sub-sequences.

### Discussion

The Mac-dBG permits to encode the color sets in C into smaller space compared to the original c-dBG and without compromising the efficiency of the ColorSet(*x*) query, for the following reasons.

1. If 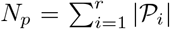 is the total number of partial color sets, then each meta color (*i, j*) can be indicated with just log_2_(*N*_*p*_) bits. Potentially long patterns, shared between several color sets, are therefore encoded once in 𝒫 and only referenced with log_2_(*N*_*p*_) bits instead of redundantly replicating their representation. Since partial color sets are encoded once, the total number of integers in 𝒫 is *at most* that in the original 𝒞. In practice, 𝒫 is expected to contain a much smaller number of integers than 𝒞.
2. Each partial color set *P*_*ij*_ can be encoded more succinctly because the permutation *π* guarantees that it only comprises integers lower-bounded by *B*_*i*−1_+1 and upper-bounded by *B*_*i*_. Hence only log_2_(*B*_*i*_ − *B*_*i*−1_) bits per integer are sufficient.
3. It is efficient to recover the original color set *C*_*t*_ from the meta color set *M*_*t*_: for each meta color (*i, j*) ∈ *M*_*t*_, sum *B*_*i*−1_ back to each decoded integer of *P*_*ij*_. Hence, we decode *strictly increasing* integers. This is, again, a direct consequence of having permuted the reference identifiers with *π*. Observe that, in principle, the representation of the color sets with partial/meta color sets could be described *without* any permutation *π* — however, one would sacrifice space (for reason 2. above) *and* query time since decoding a set would eventually need to sort the decoded integers. In conclusion, permuting the reference identifiers with *π* is an extra degree of freedom that we can exploit to improve index space and preserve query efficiency, noting that the correctness of the index is not compromised when reference identifiers are re-assigned globally.

#### The optimization problem

As already noted in Section 4.1 for the Dfc-dBG, also the effectiveness of the Mac-dBG crucially depends on the choice of the partition 𝒩 and upon the order of the references within each partition as given by the permutation *π*. There is, in fact, an evident friction between the encoding costs of the partial and meta color sets. Let *N*_*m*_ and 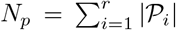 be the number of meta colors and partial color sets, respectively. Since each meta color can be indicated with log_2_(*N*_*p*_) bits, the meta color sets cost *N*_*m*_ log_2_(*N*_*p*_) bits overall. Instead, let Cost(*P*_*ij*_, *π*) be the cost (in bits) of the partial color set *P*_*ij*_ according to some encoding method. On one hand, we would like to select a large value of *r* so that *N*_*p*_ diminishes since each color set is partitioned into several, small, partial color sets, thereby increasing the chances that each partition has many repeated patterns. This will help in reducing the encoding cost for the partial color sets, i.e., the Quantity 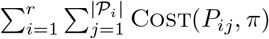. On the other hand, a large value of *r* will yield larger meta color sets, i.e., will increase *N*_*m*_. This, in turn, could erode the benefit of encoding shared patterns and would require more time to read the meta color sets. The minimum-cost partition arrangement (MPA) problem for Mac-dBG is therefore as follows.

##### Problem 3

(MPA for Mac-dBG)

Let *G*(𝒰, *𝒞*) be the compacted c-dBG built from the reference collection R = {*ℛ*_1_, … *R*_*N*_ }. Determine the partition 𝒩 = {𝒩_1_, …, *𝒩*_*r*_} of [*N*] for some *r* ≥ 1 and permutation *π* : [*N*] → [*N*] such that 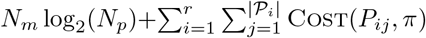 is minimum.

Depending upon the chosen encoding, smaller values of Cost(*P*_*ij*_, *π*) may be obtained when the gaps between subsequent reference identifiers are minimized. Finding the permutation *π* that minimizes the gaps between the identifiers over all partial color sets is an instance of the *bipartite minimum logarithmic arrangement* (BIMLOGA) problem as introduced by Dhulipala et al. [18] for the purpose of minimizing the cost of delta-encoded lists in inverted indexes. The BIMLOGA problem is NP-hard [18]. We note that BIMLOGA is a special case of MPA: that for *r* = 1 (one partition only) and Cost(*P*_*ij*_, *π*) being the sum of the log_2_ of the gaps between consecutive integers in the permuted set *P*_*ij*_. It follows that also MPA is NP-hard under these constraints. This result immediately suggests that it is unlikely that polynomial-time algorithms exist for solving the MPA problem for the Mac-dBG.

#### 4.3 Comparing and combining the two representations

Sections 4.1 and 4.2 introduce two solutions to the same problem; that of representing a collection of sorted integer sets taking into account for patterns of integers that repeat across the collection. These two solutions are very different, given that are based on orthogonal partitioning paradigms. As it is not therefore immediately obvious which solution may yield better results, we discuss them comparatively:

– The representative/differential color set approach can capture patterns that are formed by colors not necessarily appearing in consecutive positions, whereas the approach via partial/meta color sets necessarily needs to permute the reference identifiers to be effective. The use of permuted reference identifiers is, however, beneficial for compression as it typically reduces the number of bits to encode the differences between consecutive colors in a set.
– Both methods incur one extra level of indirection when decoding a color set *C*_*i*_ comparedto a representation that encodes each set individually. In particular, up to *r* partial color sets have to be accessed and decoded from their respective partitions; a representative and differential color set must be scanned simultaneously to decode *C*_*i*_. Recall, however, that while exactly |*C*_*i*_| integers are decoded to reconstruct *C*_*i*_ using the partial/meta color set representation, the encoding with representative/differential can instead decode more than |*C*_*i*_| integers.

The two observations above suggest that the representation via meta/partial color sets has a net advantage over that based on representative/differential color sets. However, we argue in the following that an even improved representation can be achieved when these two paradigms are *combined* together. In fact, note that both the set Δ of differential color sets and the set 𝒫 of partial color sets are, in turn, collections of sorted integer sets. Applying the same encoding strategy *recursively* would therefore seem the most straightforward option to consider. A recursive encoding makes little sense, however, as the hypothetical benefits arising from finer partitioning should have been obtained during the “outer” and only partitioning step. We therefore consider the two scenarios where: (1) the set Δ is encoded with meta/partial color sets, and (2) the set 𝒫 is encoded with representative/differential color sets. Albeit possible, the former combination is not promising as the differential color sets in each partition are expected to be very different from each other rather than similar because shared patterns are captured and encoded in the representative color sets only. (In other words, if a shared pattern emerged in the differential color sets, then this pattern would have been included in the representative color set, thus reducing the “similarity” between the differential color sets.) This is also apparent from the small example from Figure 4. The latter combination, instead, retains good potential as the partial color sets in a partition tend to be similar as well. Consider the example in Figure 5.

In conclusion, we will also experiment in Section 6 with a combined representation where the partial color sets are further compressed via representative/differential color sets.

## 5 The SCPO framework

In this section we illustrate a framework to build the compressed data structures described in Section 4, the Dfc-dBG and the Mac-dBG. We then detail the specific instance of the framework used for the experiments in this work.

As evident from their definitions, the crux of both data structures is how to perform partitioning in an effective and efficient way. We recall that the input of partitioning is different for the two data structures, i.e., the colors in C for the Dfc-dBG and the references in ℛ for the Mac-dBG. We therefore generically refer to the elements of the input as “objects” in the following. The framework is a heuristic for the introduced MPA optimization problems (Problem 2 and 3) and is based on the intuition that *similar* objects should be grouped together in the same partition so as to *increase the likeliness of having larger shared patterns*. It consists in the following four steps: (1) *Sketching*, (2) *Clustering*, (3) *Partitioning*, and (4) *Ordering*. Hence, we call it SCPO framework.

### 1. Sketching

As a pre-processing step for the actual partitioning (step 2. below), we first compute *sketches* of the objects to be partitioned. This makes the partitioning less-memory intensive and faster as it operates on smaller objects (the sketches). Furthermore, the sketches should preserve the essential information of the original objects, thus if two sketches are similar then the original objects are similar as well.

For the Dfc-dBG, we simply build one sketch for each color set. The sketches for the Mac-dBG, instead, are obtained as follows. Recall from Property 1 (Section 2) that each reference *R*_*i*_ ∈ ℛ can be spelled by a proper concatenation (a “tiling”) of the unitigs of the underlying compacted dBG. If these unitigs are assigned unique identifiers by the chosen dictionary data structure (e.g., SSHash [42,43]), it follows that each *R*_*i*_ can be seen as a list of unitig identifiers. These lists of unitig identifiers are obviously much shorter and take less space than the original DNA references. We compute one sketch for each such integer list.

### 2. Clustering

The sketches are fed as input of a clustering algorithm.

### 3. Partitioning

Once the clustering is done, each object in the input is labeled with the cluster label of the corresponding sketch so that the partition 𝒩 = {𝒩_1_, …, *𝒩*_*r*_} is uniquely determined.

### 4. Ordering

Lastly, one may *order* the objects in each partition. While this might not be relevant for the Dfc-dBG, it is definitely for the Mac-dBG because reference identifiers can be permuted to reduce the encoding cost of the lists being represented (i.e., the partial colors for the Mac-dBG). In fact, while the goal of partitioning is to factorize the color sets in their repetitive patterns, the goal of the ordering step is to assign nearby identifiers to colors that tend to co-occur in the color sets. This would mean to determine a permutation *π* that globally re-assign identifiers to references. As discussed in Section 4.2, this problem was shown to be NP-hard but good heuristics exist [18].

#### Specific framework instance

For the experiments in this work, we use the following specific instance of the framework. We build *hyper-log-log* [22] sketches of *W* = 2^10^ bytes each. As a clustering algorithm, we use a *divisive K*-means approach that does not need an *a priori* number of clusters to be supplied as input. At the beginning of the algorithm, the whole input forms a single cluster that is recursively split into two clusters until the mean squared error (MSE) between the sketches in the cluster and the cluster’s centroid is not below a prescribed threshold (which we fix to 10% of the MSE at the start of the algorithm). Let *r* be the number of found clusters. The complexity of the algorithm depends on the topology of the binary tree representing the hierarchy of splits performed. Let *Z* be the number of sketches to cluster. We recall that for the Dfc-dBG we have *Z* = *z* (the number of distinct color sets) whereas *Z* = *N* for the Mac-dBG (the number of references). In the worst case, the topology is completely unbalanced and the complexity is *O*(*WZr*); in the best case, the topology is perfectly balanced instead, for a cost of *O*(*WZ* log *r*). Note that the worst-case bound is very pessimistic because, in practice, the formed clusters tend to be reasonably well-balanced in size. Usually *z* ≫ *N*, hence we expect the clustering step performed for the Dfc-dBG to take more time and space in practice compared to that for the Mac-dBG.

In the current version of the work, we did not perform any ordering of the references. We leave the investigation of this opportunity as future work.

Lastly, in this section, it is worth noting that the approach we describe here for constructing our new compressed c-dBGs bears a conceptual resemblance to the phylogenetic compression framework recently introduced by Břinda et al. [15]. At a high level, this owes to the fact that both approaches take advantage of well-known concepts in compression and information retrieval — namely that clustering and reordering are practical and effective heuristics for boosting compression. However, while the approach by Břinda et al. focuses on clustering references so as to improve the construction of collections of disparate dictionaries, we strictly focus on the effectiveness and efficiency of the index. As such, our approach adopts a single *k*-mer dictionary and instead induces a logical partitioning over the color sets. This layout allows to avoid having to record *k*-mers that appear in multiple partitions more than once. As a result, while the phylogenetic compression framework aims to scale to immense and highly-diverse collections of references, it anticipates a primarily disk-based index in which partitions are loaded, decompressed, and searched for matches, similarly to a database search (or similarly to BLAST [8]). On the other hand, the approaches we present here place a premium on query time, and aim to enable *in-memory indexing* with interactive lookups for the purpose of fast read-mapping against the index.

## 6. Experiments

In this section, we illustrate the results of experiments conducted to assess the performance of the proposed c-dBG indexes, in comparison to the state-of-the-art methods reviewed in Section 3. We fix the *k*-mer length to *k* = 31. All experiments were run on a machine equipped with Intel Xeon Platinum 8276L CPUs (clocked at 2.20GHz), 500 GB of RAM, and running Linux 4.15.0.

**Datasets**. We perform experiments using the following pangenomes of bacteria: 3,682 *E. Coli* (EC) genomes from NCBI [1]; different collections of *S. Enterica* (SE) genomes from the “661K” collection by Blackwell et al. [11]. Specifically, we use collections of different sizes, ranging from 5,000 to 150,000 genomes. We also test a much more diverse collection of 30,691 genomes assembled from human gut samples (GB), originally published by Hiseni et al. [23]. Table 1 reports some basic statistics about these collections.

**Table 1:** Basic statistics for the tested collections, for *k* = 31.

|  | *E. Coli*<br>(EC) | *S. Enterica*<br>(SE) |  |  |  |  | Gut bacteria<br>(GB) |
| --- | --- | --- | --- | --- | --- | --- | --- |
| Genomes | 3,682 | 5,000 | 10,000 | 50,000 | 100,000 | 150,000 | 30,691 |
| Distinct color sets ($\times 10^6$) | 5.59 | 2.69 | 4.24 | 13.92 | 19.36 | 23.61 | 227.80 |
| Integers in color sets ($\times 10^9$) | 5.74 | 5.77 | 15.68 | 133.49 | 303.53 | 490.04 | 10.04 |
| Avg. color set size ($\times 10^3$) | 1.03 | 2.14 | 3.70 | 9.69 | 15.68 | 20.76 | 0.04 |
| $k$-mers in dBG ($\times 10^6$) | 170.65 | 104.69 | 239.88 | 806.23 | 1,018.69 | 1,194.44 | 13,936.86 |
| Unitigs in dBG ($\times 10^6$) | 9.31 | 4.95 | 8.24 | 30.64 | 41.16 | 49.60 | 566.39 |

Note how for the GB dataset, being much more diverse, the average color set size (computed as the ratio between the number of integers in the color sets and the number of distinct color sets) is just ≈ 44 integers — two order of magnitude smaller than that of the other collections evaluated here.

**Implementation details**.Our methods from Section 4 are not bound to a specific compression scheme nor a specific dictionary data structure, allowing one to obtain a spectrum of different space/time trade-offs depending on choices made. Here, we use these new methods to compress the color sets of the Fulgor index [21] — the state-of-the-art c-dBG index and our own previous work (see Section 3). Like in Fulgor, we therefore use the SSHash data structure [42,43] to represent the set of unitigs 𝒰 and the same mechanism to map unitigs to their color sets (see Fact 1). Very importantly, note that these choices directly imply that our new indexes fully exploit the unitig properties described in Section 2 as Fulgor does. We therefore experiment with the following indexes: the differential-colored version of Fulgor, or “d-Fulgor” in the following; the meta-colored version, or “m-Fulgor”; and the “md-Fulgor” representation, obtained by encoding with representative/differential color sets each partial color set as explained in Section 4.3.

Both representative and differential color sets in d-Fulgor are simply encoded by taking the gaps between consecutive integers and representing each gap with Elias’ δ code. For the partial color sets of m-Fulgor, we adopt the same compression methods as used to represent the color sets in Fulgor (see the description at the end of Section 3). Each meta color list is instead a list of log_2_(*N*_*p*_)-bit integers, *N*_*p*_ *>* 0 being the total number of partial color sets.

For the md-Fulgor index — our most succinct variant of Fulgor — we also use a different schemes for the meta colors to improve compression further. Recall from Section 4.2 that each meta color is an integer pair (*i, j*), where *i* indicates a partition and *j* indicates the offset of the partial color set *P*_*ij*_. A list of meta colors *M*_*t*_ = [(*i*_1_, *j*_1_), (*i*_2_, *j*_2_), …] can therefore be decomposed into the two integer lists [*i*_1_, *i*_2_, …] and [*j*_1_, *j*_2_,], which we referred to as *first* and *second* components in the following. For example, the meta color list *M*_5_ = [(1, 4), (2, 3), (3, 2), (4, 4)] from Figure 5 can be decomposed into its first component [1, 2, 3, 4] and its second component [4, 3, 2, 4]. Note that while the first component is always a sorted list of all elements are distinct, this is not true in general for the second component. We thus use different encoding schemes for first and second components.

First, observe that — especially when the number of partitions is small — we expect a small number of *distinct* first components, say *f*, compared to the total number of color sets, *z*. Hence, it is convenient to keep the distinct first components in an array *A*[1..*f*] and, for each meta color, specify the index of the array’s entry corresponding to the first component, using log_2_(*f*) bits. Even better, we can sort the meta color lists by first component and implement the succinct map via Rank explained in Fact 1. Thus, we use a bitvector *b*[1..*z*] plus the array *A*[1..*f*] and recover the first component of the *t*-th meta color list as *A*[*p*] where *p* = Rank_1_(*F, t*) + 1.

The second components, instead, are not so repetitive and, as noted above, the integers are not necessarily distinct and increasing. Thus, we use the following variable-length encoding: given the meta color (*i, j*), we encode the integer *j* using log_2_(|𝒫_*i*_|) bits. For the example meta color list *M*_5_ we have used above, we will encode its second component [4, 3, 2, 4] using log_2_(|𝒫_1_|) + log_2_(|𝒫_2_|) + log_2_(|𝒫_3_|) + log_2_(|𝒫_4_|) bits, that is log_2_(5) + log_2_(3) + log_2_(2) + log_2_(6) = 3 + 2 + 1 + 3 = 9 bits.

**Compared indexes**. Throughout this section, we compare our c-dBG data structures — d-Fulgor, m-Fulgor, and md-Fulgor — against the following indexes: the original Fulgor index [21], Themisto [3], MetaGraph [27,28,29], and COBS [10]. Links to the corresponding software libraries can be found in the References. We use the `C++` implementations from the respective authors. All software (including ours) was compiled with `gcc 11.1.0`.

We provide some details on the tested tools. Both Themisto and COBS were built under default parameters as suggested by the authors, that is: with option `-d 20` for Themisto for better space effectiveness; in COBS, we have shards of at most 1024 references where each Bloom filter has a false positive rate of 0.3 and one hash function to accelerate lookup operations. MetaGraph indexes were built with the *relaxed row-diff* BRWT data structure [28] using a workflow available at https://github.com/theJasonFan/metagraph-workflows that we wrote with the input of the MetaGraph authors.

### 6.1 Space effectiveness

Table 2 reports the total *on disk* index size for all of the methods evaluated. In particular, Table 2a illustrates the effectiveness of the methods introduced in Section 4. Compared to the Fulgor index that was previously shown to achieve the most desirable space/time trade-off [21], our three methods, in order from left to right in the table, offer a progressive reduction is space. The improvement in the representation of the color sets is even more dramatic as the number of references in the dataset increases. In fact, as the size of the collection grows and more repetitive patterns in the color sets appear, our repetition-aware compression algorithms are able to better capture and reduce this redundancy. To make a concrete example, on the larger SE-150K dataset, the space spent to represent the color sets goes from 68.61 GB in Fulgor to:

**Table 2:** Index space in GB, broken down by space required for indexing the *k*-mers in the dBG (SSHash for all Fulgor variants which we hence report only once, SBWT for Themisto, and BOSS for MetaGraph) and data structures required to encode color sets and map *k*-mers to the sets. In (a), we also indicate in parentheses the percentage of space taken by the color sets with respect to the total. (For COBS, we just report the total index size which coincides with the space of the approximate color sets.)

| Dataset | Fulgor |  |  | d-Fulgor |  | m-Fulgor |  | md-Fulgor |  |
| --- | --- | --- | --- | --- | --- | --- | --- | --- | --- |
|  | dBG | Color sets | Total | Color sets | Total | Color sets | Total | Color sets | Total |
| EC | 0.29 | 1.36 (83%) | 1.65 | 0.45 (61%) | 0.74 | 0.40 (58%) | 0.69 | 0.24 (45%) | 0.52 |
| SE-5K | 0.16 | 0.59 (79%) | 0.75 | 0.20 (56%) | 0.36 | 0.16 (50%) | 0.32 | 0.11 (40%) | 0.27 |
| SE-10K | 0.35 | 1.66 (83%) | 2.01 | 0.48 (58%) | 0.83 | 0.34 (49%) | 0.70 | 0.22 (39%) | 0.57 |
| SE-50K | 1.25 | 17.03 (93%) | 18.29 | 4.31 (77%) | 5.57 | 2.08 (62%) | 3.34 | 1.38 (52%) | 2.64 |
| SE-100K | 1.71 | 40.71 (96%) | 42.43 | 9.37 (84%) | 11.10 | 3.75 (68%) | 5.47 | 2.26 (57%) | 3.98 |
| SE-150K | 2.02 | 68.61 (97%) | 70.65 | 15.73 (89%) | 17.77 | 5.27 (72%) | 7.31 | 3.22 (61%) | 5.26 |
| GB | 21.29 | 15.54 (42%) | 36.83 | 7.51 (26%) | 28.81 | 9.16 (30%) | 30.46 | 6.19 (23%) | 27.48 |

(b)
| Dataset | Themisto |  |  | MetaGraph |  |  | COBS |
| --- | --- | --- | --- | --- | --- | --- | --- |
|  | dBG | Color sets | Total | dBG | Color sets | Total | Total |
| EC | 0.22 | 1.85 | 2.08 | 0.10 | 0.23 | 0.33 | 7.53 |
| SE-5K | 0.14 | 1.29 | 1.43 | 0.07 | 0.19 | 0.26 | 9.11 |
| SE-10K | 0.32 | 3.50 | 3.81 | 0.13 | 0.38 | 0.51 | 18.68 |
| SE-50K | 1.07 | 32.42 | 33.48 | 0.36 | 1.95 | 2.31 | 88.61 |
| SE-100K | 1.35 | 75.94 | 77.28 | 0.45 | 3.50 | 3.95 | 173.58 |
| SE-150K | 1.58 | 125.16 | 126.74 | NA | NA | NA | 265.49 |
| GB | 18.33 | 30.88 | 49.21 | 5.23 | 4.77 | 10.00 | 21.23 |

– 15.73 GB in d-Fulgor (4.3× smaller space),
– 5.27 GB in m-Fulgor (13× smaller space),
– 3.22 GB in md-Fulgor, for a total reduction in space of more than 21.3×.

We also report in the table the percentage of space taken by the color sets relative to the total size of the indexes (in gray shade). For the largest index compared to here, the original Fulgor, the percentage illustrates how the space for the color sets tends to dominate the whole representation space for c-dBGs. However, note how the percentage gradually diminishes moving from the left to the right side of the table, i.e., as compression gradually improves. In some instances, the reduction offered by our algorithms surprisingly makes SSHash a “heavy” component of the index (which takes a constant fraction of the total space across all Fulgor variants).

Remarkably, observe that d-Fulgor compresses better than m-Fulgor the color sets for the GB pangenome (7.51 GB vs. 9.16 GB). This suggests that the method based on representative/differential color sets works better than the other one when the sets being represented are small.

We now turn our attention to the comparison against the other state-of-the-art methods evaluated in Table 2b. Note that the original Fulgor index already improves over the space usage of Themisto, thus making even more apparent the reduction in space of our newly introduced variants compared to Themisto. In fact, the only index whose size is competitive to ours is MetaGraph in the relaxed row-diff BRWT configuration — at least in the cases where we were able to construct the latter within the construction resource constraints. However, as we observe in Section 6.2, unlike the other indexes evaluated, the *on disk* index size MetaGraph is not representative of the working memory required for query when using the (recommended and default) “batch” mode query. The color sets of md-Fulgor also mostly require less space than those of MetaGraph.

The COBS index, despite being approximate, is consistently and considerably larger than all of the other (exact) indexes, except for the the gut bacteria collection (`GB`). The differing behavior on `GB` likely derives from the fact that the diversity of that data cause the exact indexes to spend a considerable fraction of their total size on the representation of the *k*-mer dictionary itself (e.g., 18 − 21.3 GB). However COBS, by design, eliminates this component of the index entirely.

### 6.2 Query efficiency

Table 3 reports the query times of the indexes on a high-hit workload, e.g., when (more than) 90% of the queries have a non-empty result. (Performance on a low-hit workload is less informative since few *k*-mers would actually be found in the indexes, thus testing the speed of negative *k*-mer lookups against the dictionary data structure, rather than the time spent in processing the color sets.) The time we measure for this experiment is that for performing *pseudoalignment*. There are several pseudoalignment algorithms (see [21, Section 4] for an overview) that standard c-dBG data structures directly support; here we focus on the *full intersection* algorithm (Alg. 1 from [21]). Given a query string *Q*, we consider it as a set of *k*-mers. Let 𝒦 (*Q*) = {*x* ∈ *Q*|ColorSet(*x*)≠ ∅}. The full intersection method computes the intersection between the color sets of all the *k*-mers in 𝒦 (*Q*).

**Table 3:**
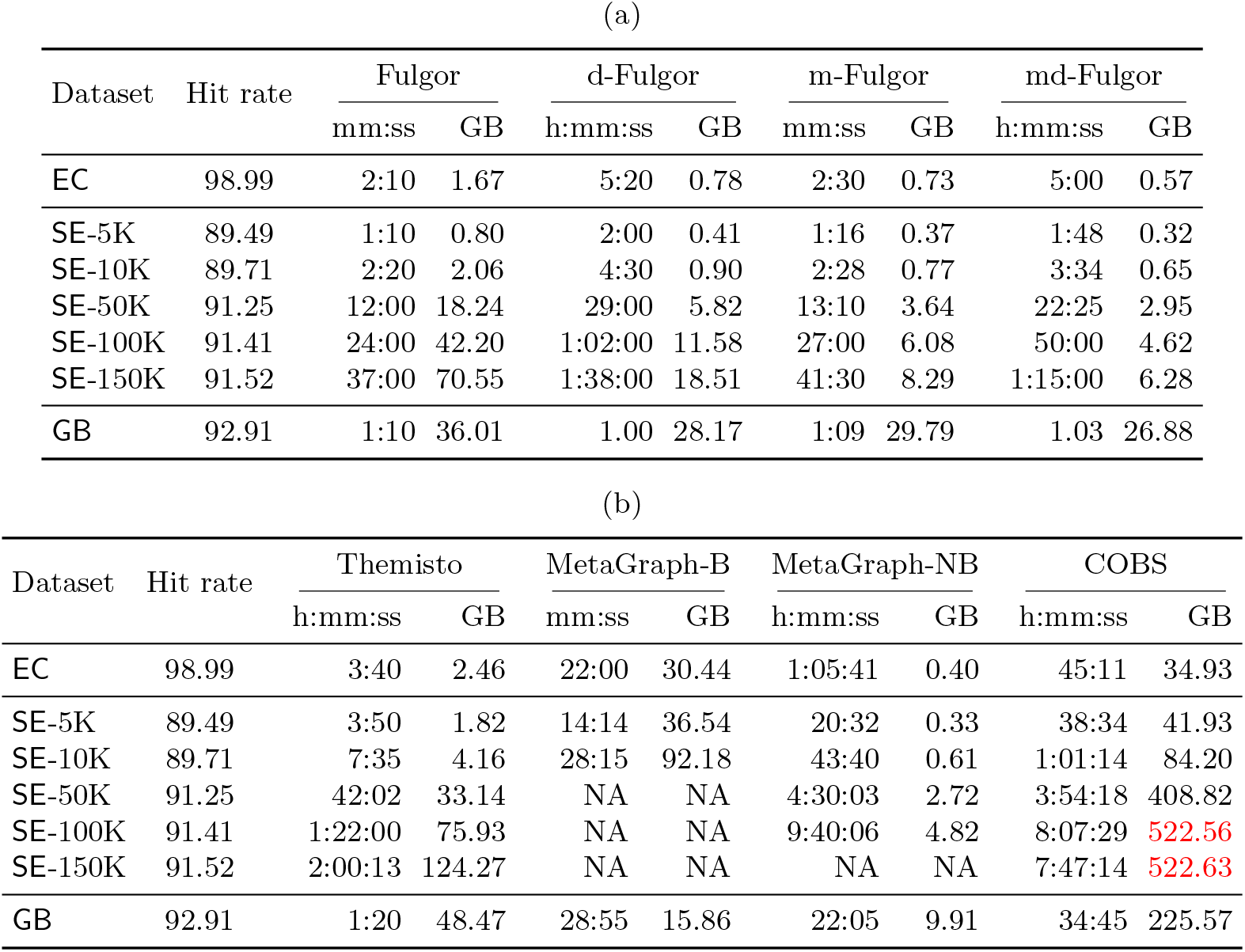
Total query time (elapsed time) and memory used during query (max. RSS) as reported by /`usr/bin/time-v`, using 16 processing threads. The read-mapping output is written to `/dev/null` for this experiment. We also report the mapping rate in percentage (fraction of mapped read over the total number of queried reads). The query algorithm used here is full-intersection. The “B” query mode of MetaGraph corresponds to the batch mode (with default batch size); the “NB” corresponds to the non-batch query mode instead. In red font we highlight the workloads exceeding the available memory (*>* 500 GB).

The queried reads consist of all FASTQ records in the first read file of the following accessions: `SRR1928200` for `EC, SRR801268` for `SE`, and `ERR321482` for `GB`. These files contain several million reads each. Timings are relative to a second run of each experiment, where the indexes are loaded from the disk cache (which benefits the larger indexes more than the smaller ones).

Consistent with previously reported results [21], we find that among existing indexes, Fulgor provides the fastest queries. The d-Fulgor variant results in being approximately 2× slower than Fulgor but also offers much better compression effectiveness. The slowdown is due to the differential color sets taking more time to be decoded than the original color sets (i.e., linear time in the *sum* of the lengths of the representative and the differential sets).

Vertical partitioning, instead, opens the possibility to achieve even faster query times than a traditional c-dBG, due to the manner in which the partitions factorize the space of references, if a *two-level* intersection algorithm is employed for pseudoalignment. First, only meta color sets are intersected (thus, without any need to access the partial color sets) to determine the partitions in common to all color sets being intersected. Then, only the common partitions are considered. Two cases can happen for each partition. Case (1): the meta color is the same for all color sets being intersected. In this case, the result of the intersection is *implicit* and it suffices to decode the partial color set indicated by the meta color. Case (2): the meta color is not the same, hence we have to compute the intersection between different partial color sets. This optimization is clearly beneficial when the color sets being intersected have very few partitions in common, or when they have identical meta color sets. This is the reason why m-Fulgor does not not sacrifice query efficiency compared to Fulgor, as expected, despite the significant reduction in space.

All Fulgor variants are instead equally efficient when indexing the `GB` dataset. The reason is that, on average, very small color sets are being decoded and intersected.

Taking also into account the space of the indexes as discussed in Section 6.1, two main conclusions emerge:

– The m-Fulgor variant dominates completely the original Fulgor index, as it is considerably more space efficient and equally fast to query.
– The *combination* of vertical and horizontal partitioning in the md-Fulgor index, partially mitigates the slowdown given by d-Fulgor. Overall, this combination makes the md-Fulgor more than one order of magnitude smaller than the original Fulgor index with a slowdown in query processing of less than 2×. We consider this space/time trade-off very reasonable for the sake of indexing larger and larger c-dBGs in internal memory. As we note below, md-Fulgor is still 2× faster than Themisto, the next fastest index in the literature.

After Fulgor and m-Fulgor, we note that Themisto is the next fastest index, followed by MetaGraph in batch query mode. Our most succinct but also slower version, the md-Fulgor index, is still roughly 2× faster than Themisto. The query speeds of COBS and of MetaGraph when not executed in batch mode are much lower than that of the other indexes, in some cases being (more than) an order of magnitude slower.

Critically, as anticipated in Section 6.1, it is not the case with all indexes evaluated here that the size of the index *on disk* is a good proxy for the memory required to actually query the index. Specifically, for MetaGraph, when used in batch query mode (“B”), the required memory can exceed the on-disk index size by up to 2 orders of magnitude, and in several tests this resulted in the exhaustion of available memory and an inability to complete the queries under the tested configuration. On the other hand, all Fulgor variants, Themisto, and MetaGraph when not executed in batch mode (“NB”) require only a small constant amount of working memory beyond the size of the index present on disk.

The COBS query is generally much slower than the other indexes, except for MetaGraph in non-batch mode. This is likely because COBS partitions the input collection into shards of references of roughly the same size prior to indexing. This permits to build Bloom filters of different sizes: filters belonging to different shards have a different number of bits allocated, hence saving space compared to the case where all references are represented with filters of the same size. At query time, however, a *k*-mer lookup has to be resolved by every shard and individual results combined.

### 6.3 Construction time and space

In Table 4 we consider the resources needed to build the indexes. Both d-Fulgor and m-Fulgor are built from a Fulgor index to which we apply the SCPO framework (Section 5). Similarly, the md-Fulgor is built from an m-Fulgor index. For this reason, the time reported in Table 4 for these indexes has to be summed to the time needed to first build another variant of Fulgor. The SCPO framework is single-threaded except for the construction and clustering of the sketches.

**Table 4:** Total index construction time (elapsed time) and GB of memory used during construction (max. RSS), as reported by `/usr/bin/time -v`, using 48 processing threads. The reported time includes the time to serialize the index on disk. In red font we highlight the constructions exceeding the available memory (*>* 500 GB) and for which we had to cap the maximum memory usage to 100 GB. The time for the three Fulgor variants is that for running the SCPO construction, so it has to be summed to the time needed to first build an index to partition: the time of both d-Fulgor and m-Fulgor must be summed to that of Fulgor, and the time for md-Fulgor must be summed to that of m-Fulgor.

| Dataset | Fulgor |  | d-Fulgor |  | m-Fulgor |  | md-Fulgor |  |
| --- | --- | --- | --- | --- | --- | --- | --- | --- |
|  | h:mm | GB | h:mm | GB | h:mm | GB | h:mm | GB |
| EC | 0:06 | 17 | +0:12 | 11 | +0:05 | 3 | +0:14 | 4 |
| SE-5K | 0:04 | 13 | +0:06 | 8 | +0:04 | 1 | +0:05 | 3 |
| SE-10K | 0:09 | 24 | +0:09 | 14 | +0:10 | 3 | +0:11 | 4 |
| SE-50K | 1:13 | 44 | +0:43 | 105 | +1:50 | 22 | +0:30 | 13 |
| SE-100K | 2:56 | 74 | +1:20 | 207 | +4:37 | 48 | +0:43 | 20 |
| SE-150K | 4:36 | 137 | +1:55 | 305 | +7:41 | 77 | +0:55 | 25 |
| GB | 2:27 | 115 | +10:00 | 182 | +0:31 | 69 | +7:12 | 127 |

(b)
| Dataset | Themisto |  | MetaGraph |  | COBS |  |
| --- | --- | --- | --- | --- | --- | --- |
|  | h:mm | GB | h:mm | GB | h:mm | GB |
| EC | 0:19 | 17 | 0:46 | 149 | 0:03 | 6 |
| SE-5K | 0:11 | 13 | 0:47 | 191 | 0:09 | 8 |
| SE-10K | 0:25 | 24 | 1:50 | 219 | 0:17 | 16 |
| SE-50K | 2:32 | 96 | 14:16 | **120** | 1:41 | 82 |
| SE-100K | 6:25 | 202 | 26:40 | **104** | 2:37 | 84 |
| SE-150K | 10:00 | 323 | NA | NA | 4:54 | 159 |
| GB | 6:21 | 184 | 10:50 | **100** | 0:22 | 17 |

We note that the m-Fulgor index is slower to build compared to a d-Fulgor index, despite the fact that d-Fulgor re-builds a (permuted) SSHash dictionary and clusters many more sketches. The reason lies in the different parameters that we used for the clustering of the sketches. For d-Fulgor, we do not impose any constraint of the minimum size of a cluster, hence allowing the algorithm to produce small and many clusters. On the contrary, the number of clusters must be kept under control for the m-Fulgor variant as this directly impacts on the length of the meta color lists (hence, their space usage). For this reason, the clustering algorithm attempts to greedily re-assign sketches from small clusters to big clusters, with the goal of reducing the number of clusters. The memory used during the building of d-Fulgor is nonetheless much higher than that of m-Fulgor for the much larger number of sketches that are being clustered. The md-Fulgor variant can instead be built very economically from an existing m-Fulgor index.

Despite not being heavily engineered yet, the end-to-end construction of our indexes is competitive to that of Themisto and much faster than that of MetaGraph. The fastest indexes to build are Fulgor and COBS, the latter being even faster on the GB collection for reasons already explained (i.e., it does not build any exact dictionary for the *k*-mers). The tested MetaGraph configuration is significantly slower to build than all the other indexes; for example, we were unable to build the index for SE-150K within 3 days and using 48 parallel threads (the construction also produced more than 1 TB of intermediate files).

## 7 Conclusions and future work

In this work, we have introduced new compressed representations for the colored de Bruijn graph, where repetitive patterns within colors are encoded once as to improve the memory usage of pseudoalignment queries. More specifically, we have applied our compression algorithms to represent the color sets as stored in the recently introduced Fulgor index as it embodies the most favorable space vs. time trade-off, and we have introduced two distinct, and largely orthogonal, approaches for factorizing and compressing redundant patterns among the color sets. One scheme, d-Fulgor, is a “horizontal” compression method which performs a representative/differential encoding of the color sets. The other scheme, m-Fulgor, is a “vertical” compression method which instead decomposes the color sets into meta and partial color sets. Crucially, these methods exploit different characteristics for compression, and so they can be combined to achieve an even greater compression of the color sets. We show the effect of this combination with an index that we call md-Fulgor.

We perform an extensive experimental analysis across several datasets to assess these different schemes and compare them against alternative representations of the c-dBG. From our analysis, we conclude that: (1) the meta-colored version of Fulgor, m-Fulgor, does not introduce any new trade-off compared to the original Fulgor index but simply *supersedes* it, given that it is considerably smaller and equally fast; (2) the meta-differential-colored variant, md-Fulgor, is even more compact with a relatively minor query overhead compared to Fulgor, especially compared to the space savings it provides.

Our representations provide a substantially-improved new reference point for the problem of indexing c-dBGs in compressed space, as apparent from Figure 6. Our most succinct index, md-Fulgor, is competitive withe the smallest variant of MetaGraph but an order of magnitude faster to query; and up to 20× smaller than Themisto and still twice as fast. We believe this improved performance has the potential of enabling large-scale color set queries across a range of applications.

**Fig. 6:**
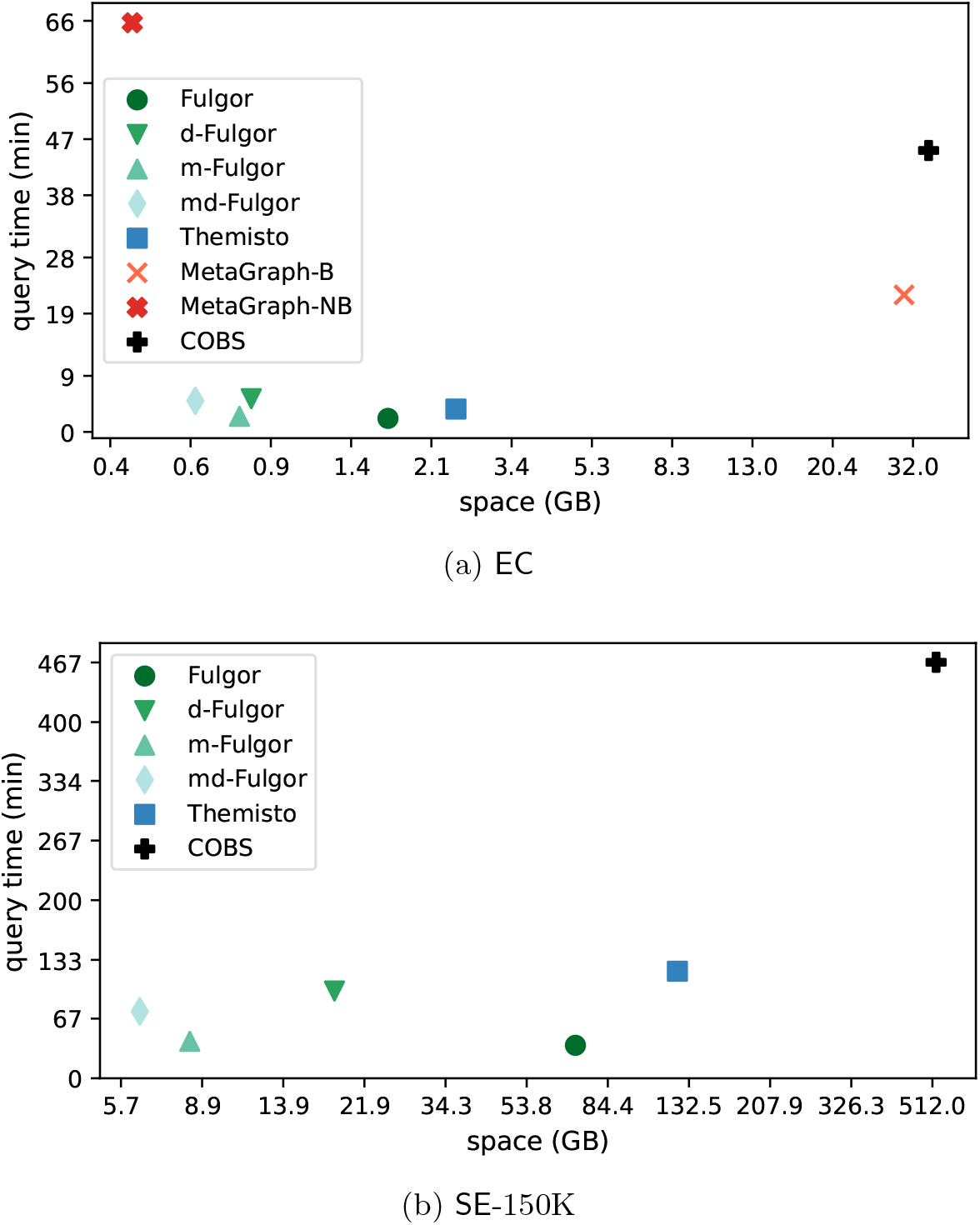
The same data from Table 3 but shown as space vs. time trade-off points, for two example datasets.

Future work will focus on different aspects of improving the index and relevant operations upon it. First, we would like to accelerate pseudoalignment queries in general, and especially for the md-Fulgor index. Second, we will provide a better engineered build pipeline for our indexes. Third, we would like to explore the effect of approximately optimal ordering within the partial color sets (i.e., the “O” step of the SCPO framework we have introduced). Lastly, we plan to extend the indexing capabilities of Fulgor by annotating its graph with more information, like *k*-mer abundances and their positions in the references.

## Fundings

This work is supported by the NIH under grant award numbers R01HG009937 to R.P.; the NSF awards CCF-1750472 and CNS-1763680 to R.P. and DGE-1840340 to J.F. Funding for this research has also been provided by the European Union’s Horizon Europe research and innovation programme (EFRA project, Grant Agreement Number 101093026). This work was also partially supported by DAIS – Ca’ Foscari University of Venice within the IRIDE program.

## Declarations

R.P. is a co-founder of Ocean Genomics inc.

## Footnotes

4 In our previous work [21,44], we use a different nomenclature and call “color” the set of references that contain a *k*-mer, so that each *k*-mer is logically labelled with a single “color” in the graph. Under this view, a color is what is called a “color class” by Pandey et al. [40]. However, here we adopt the original terminology of Iqbal et al. [25] to avoid inconsistencies or misunderstandings and to accord with the prevailing nomenclature in the related literature.

5 For ease of notation, we write log_2_(*x*) instead of ⌊log_2_(*x* − 1)⌋ + 1 for any *x* ≥ 1 and assume that log_2_(0) = 0.

## Notes

https://github.com/jermp/fulgor

### Competing Interest Statement

The authors have declared no competing interest.

